# nMAGMA: a network enhanced method for inferring risk genes from GWAS summary statistics and its application to schizophrenia

**DOI:** 10.1101/2020.08.15.250282

**Authors:** Anyi Yang, Jingqi Chen, Xing-Ming Zhao

## Abstract

**Motivation:** Annotating genetic variants from summary statistics of genome-wide association studies (GWAS) is crucial for predicting risk genes of various disorders. The multi-marker analysis of genomic annotation (MAGMA) is one of the most popular tools for this purpose, where MAGMA aggregates signals of single nucleotide polymorphisms (SNPs) to their nearby genes. However, SNPs may also affect genes in a distance, thus missed by MAGMA. Although different upgrades of MAGMA have been proposed to extend gene-wise variant annotations with more information (e.g. Hi-C or eQTL), the regulatory relationships among genes and the tissue-specificity of signals have not been taken into account.

**Results:** We propose a new approach, namely network-enhanced MAGMA (nMAGMA), for gene-wise annotation of variants from GWAS summary statistics. Compared with MAGMA and H-MAGMA, nMAGMA significantly extends the lists of genes that can be annotated to SNPs by integrating local signals, long-range regulation signals, and tissue-specific gene networks. When applied to schizophrenia, nMAGMA is able to detect more risk genes (217% more than MAGMA and 57% more than H-MAGMA) that are reasonably involved in schizophrenia compared to MAGMA and H-MAGMA. Some disease-related functions (e.g. the ATPase pathway in Cortex) tissues are also uncovered in nMAGMA but not in MAGMA or H-MAGMA. Moreover, nMAGMA provides tissue-specific risk signals, which are useful for understanding disorders with multi-tissue origins.

## Introduction

With the great power of genome-wide association studies (GWAS), a lot of risk loci have been identified for various disorders, which can help pinpoint the genetic mechanisms underlying diseases. For example, 145 risk loci have been detected from a recent GWAS study of schizophrenia (SCZ), and candidate causal genes were identified from these loci [1]. However, there are generally very few genetic variants that reach genome-wide significance in GWAS, and these variants can only explain a small fraction of disorder heritability collectively [2]. It has been found that some variants with weak effects may also contribute to disease risks [3-5], where the variants can be aggregated to a smaller group of genes based on their putative impact on gene function. Therefore, gene-wise variant annotations can help detect potential risk genes and functional convergence among massive amounts of weak-effect variants.

Recently, some computational approaches have been proposed to aggregate genetic variants with weak effects to nearby genes, among which the multi-marker analysis of genomic annotation (MAGMA) is most widely used [6]. Except for aggregating variants to genes, MAGMA has shown outstanding statistical power to control for the linkage disequilibrium information as well as other confounding factors compared with conventional gene-set analyses [7]. Nevertheless, MAGMA tends to suffer from several obvious limitations. Firstly, a user-defined ‘gene window’ is required by MAGMA for annotating SNPs to genes. The most suitable window size is an open question, where a window that is too large could increase the probability of SNPs being assigned to genes without any functional connection [8]. Secondly, MAGMA assigns an SNP to genes only based on their physical location, therefore large amounts of SNPs located in deep intergenic regions may be missed. It has been reported that intergenic SNPs can contribute to disease risks through long-range regulation [9], thus should be included to better identify disease genes. Thirdly, MAGMA does not consider tissue specificity, whereas the role of different tissues differs in their contribution to diseases, such as in the case of neuropsychiatric traits [10]. Finally, the functional relationships among genes are not considered in MAGMA, while the genes achieve certain functions by interacting with each other [11].

Several tools aiming to enhance MAGMA have been proposed to address some of the issues mentioned above. For example, H-MAGMA (Hi-C-coupled MAGMA) identified tissue-specific SNP-gene pairs by incorporating chromatin interaction profiles from brain tissues, and obtained promising results for psychiatric and neurodegenerative disorders [12]. eMAGMA (eQTL-informed method) detected risk genes based on eQTLs instead of assigning SNPs to physical-location nearby genes [13]. Despite the obvious enhancement made by H-MAGMA and eMAGMA, neither method considers the functional interactions among genes or the tissue-specificity of risk signals. It has been reported that the gene interactions derived from their co-expression profiles can help prioritize candidate genes associated with SCZ [14]. By integrating gene networks inferred from functional convergence into a Bayesian model, iRIGS inferred more credible risk genes around each independent risk loci (or index SNPs) in SCZ GWAS data [15]. Additionally, neither H-MAGMA nor eMAGMA method has considered the tissue-specificity of risk signals.

Here, we present a network-enhanced version of MAGMA, namely nMAGMA, to identify risk genes from GWAS data. In addition to the Hi-C and eQTL information used by H-MAGMA and eMAGMA, nMAGMA innovatively incorporates gene interactions derived from topological overlap matrix (TOM) [16] of weighted gene co-expression network analysis (WGCNA) [17, 18] into genetic variant annotations. Furthermore, nMAGMA is built over four tissues associated with SCZ, including Cortex, Hippocampus, Liver and Small Bowel [10, 19-21]. When applied to SCZ GWAS data, nMAGMA is able to detect more biological meaningful risk genes and gene-sets involved in schizophrenia. As such, our method may provide important insights into the biological mechanisms underlying diseases that are previously not highlighted by other methods. Additionally, the framework of nMAGMA can be flexibly expanded by adding new kinds of gene networks and omics data from any tissues, to accommodate to other traits or diseases.

## Results

### Overview of nMAGMA

Mapping SNPs to genes is one of the most important steps for post-GWAS analyses, and the main goal of our work is to assign SNPs to genes more thoroughly and accurately. Figure 1 provides a schematic view of the framework. In brief, several types of supporting evidence for the connections between individual SNPs and genes, including network-induced information, significant eQTL results, and Hi-C data, were used in the gene-wise variant annotation to generate tissue-specific annotation files for downstream gene and gene-set analysis, under the powerful framework of MAGMA. We refer to this enhanced framework as nMAGMA. After retrieving SNPs from the GWAS summary statistics for interested traits or diseases, for fundamental annotation, SNPs located in or near gene regions are assigned to their nearest genes based on their genomic coordinates. For increasing the utilization of SNPs—especially those in deep intergenic regions, additional SNP-gene pairs are generated based on chromatin interaction and significant eQTL relationships (Materials and Methods). We reason that signals captured by Hi-C and eQTL may explain a large portion of heritability which cannot be explained by simple location-based annotation. Importantly, we adopted WGCNA in order to explicitly infer relationships between genes in the biological networks. The TOM value (i.e. the corresponding value from the topological overlap matrix) between two protein-coding genes is affected not only by the direct expressional correlation between them, but also by the indirect correlation with an intermediate gene which might be noncoding, such as a lincRNA. Two genes tend to have a high TOM value when they highly interconnect with each other directly and indirectly [17, 22]. The main difference between nMAGMA and contemporary MAGMA-based strategies (i.e. H-MAGMA and eMAGMA) is that it integrates network information which is tissue-specific, resulting in annotations that are biologically and topologically more meaningful. Another advantage of nMAGMA is that it can be extended feasibly by adding new genomic signals and network information. Details of nMAGMA can be seen in Materials and Methods.

**Figure 1:**
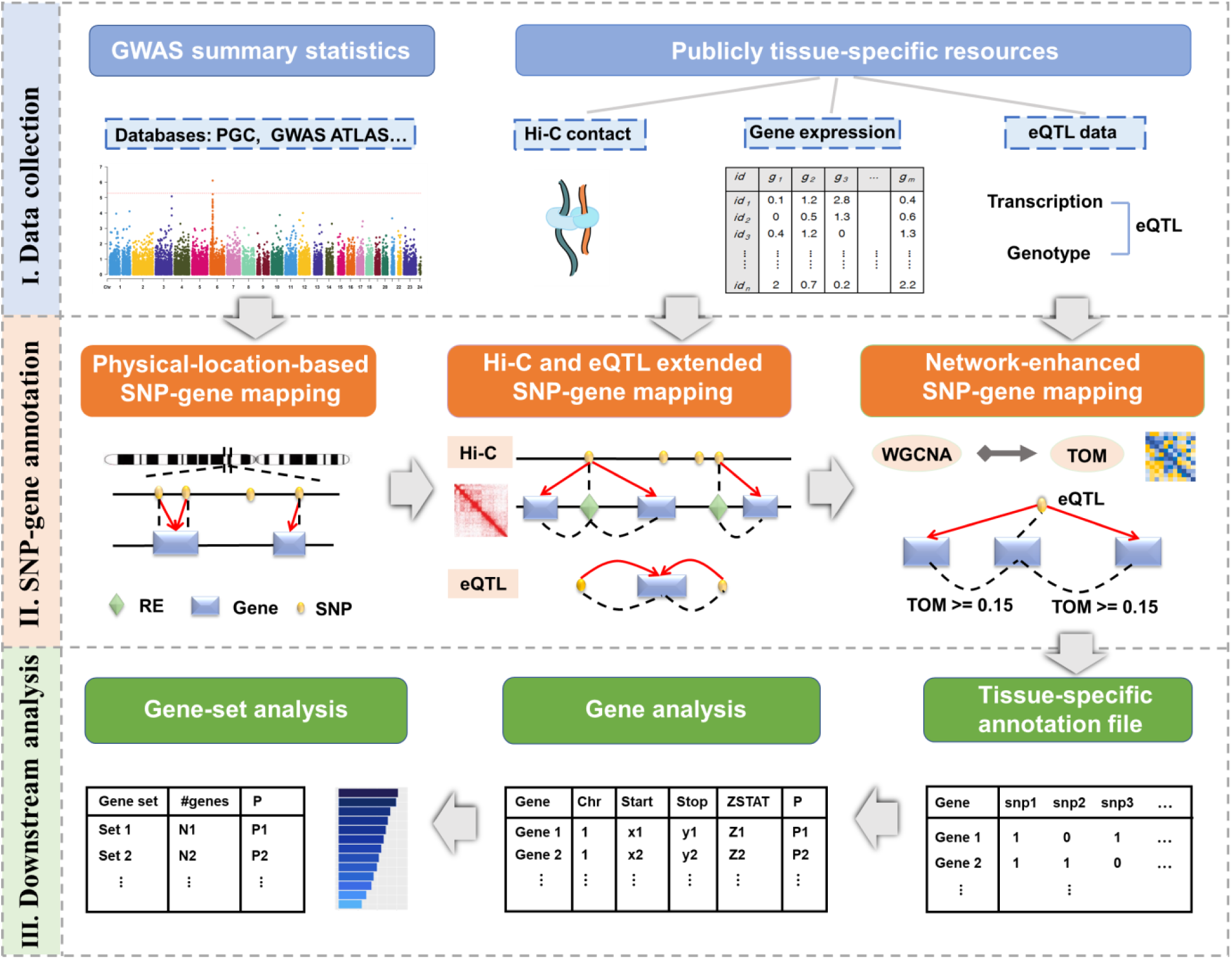
The schematic overview of the nMAGMA approach. nMAGMA uses Hi-C, eQTL, Gene regulatory interactions through TOM matrix calculated by WGCNA, as well as the physical-location-based mapping to get an expanded gene-wise annotation file of SNPs. The dashed line means there exists some relationship between the two elements, and the red arrow means the new assignment of the SNP under such relationship. The pictures representing Hi-C interaction matrix are adapted from Rao *et al*. [23].

### Application of nMAGMA to SCZ GWAS data

We applied nMAGMA to the summary statistics of two large SCZ GWAS studies of European ancestry [2, 24]. Considering the availability of Hi-C data, we only focused on the four tissues previously found related to SCZ, i.e. Cortex, Hippocampus, Liver and Small Bowel [10, 19-21]. In total, 20,635 protein-coding genes and 17,224 gene-sets were considered for further analysis, and the “SNP-wise=mean” parameter from MAGMA was adopted through all analysis.

Compared with the results by combinations of MAGMA and Hi-C as well as eQTL, nMAGMA can detect more risk genes across all four tissues as shown in Figure 2A. From the results, we can see that both Hi-C and eQTL can significantly extend the list of genes that can be mapped to the coordinates of SNPs, while the network information can further expand the number of significant genes (by 11% ∼ 25%) on top of the combination of MAGMA, Hi-C data, and eQTL data. As shown in Figure 2B, we also noticed that the median number of SNPs that can be mapped to each gene was also significantly increased by both Hi-C and eQTL compared with using only physical location information, which confirm that some remote SNPs can be mapped to genes with the help of Hi-C and eQTL. When looking at the overlap of SNPs that can be mapped to genes by Hi-C or eQTL, we found that only few SNPs (about 5%) can be detected by both datasets as shown in Figure 2C, which implies that Hi-C and eQTL can complement with each other when annotating the SNP-gene pairs, where the cis-eQTLs can annotate SNPs from the proximal regions of genes while the chromatin interaction profiles from Hi-C can help detect SNPs from regions far away from the genes. Figure 2D shows the Venn diagram of the risk genes inferred by nMAGMA across four tissues, where we can see the tissue-specificity of the risk genes detected in a certain tissue. Among the 683 genes that can be detected in all four tissues of nMAGMA, 591 of them (86.5%) can also be detected by MAGMA, suggesting that these genes may be indeed affected by SNPs inside it.

**Figure 2:**
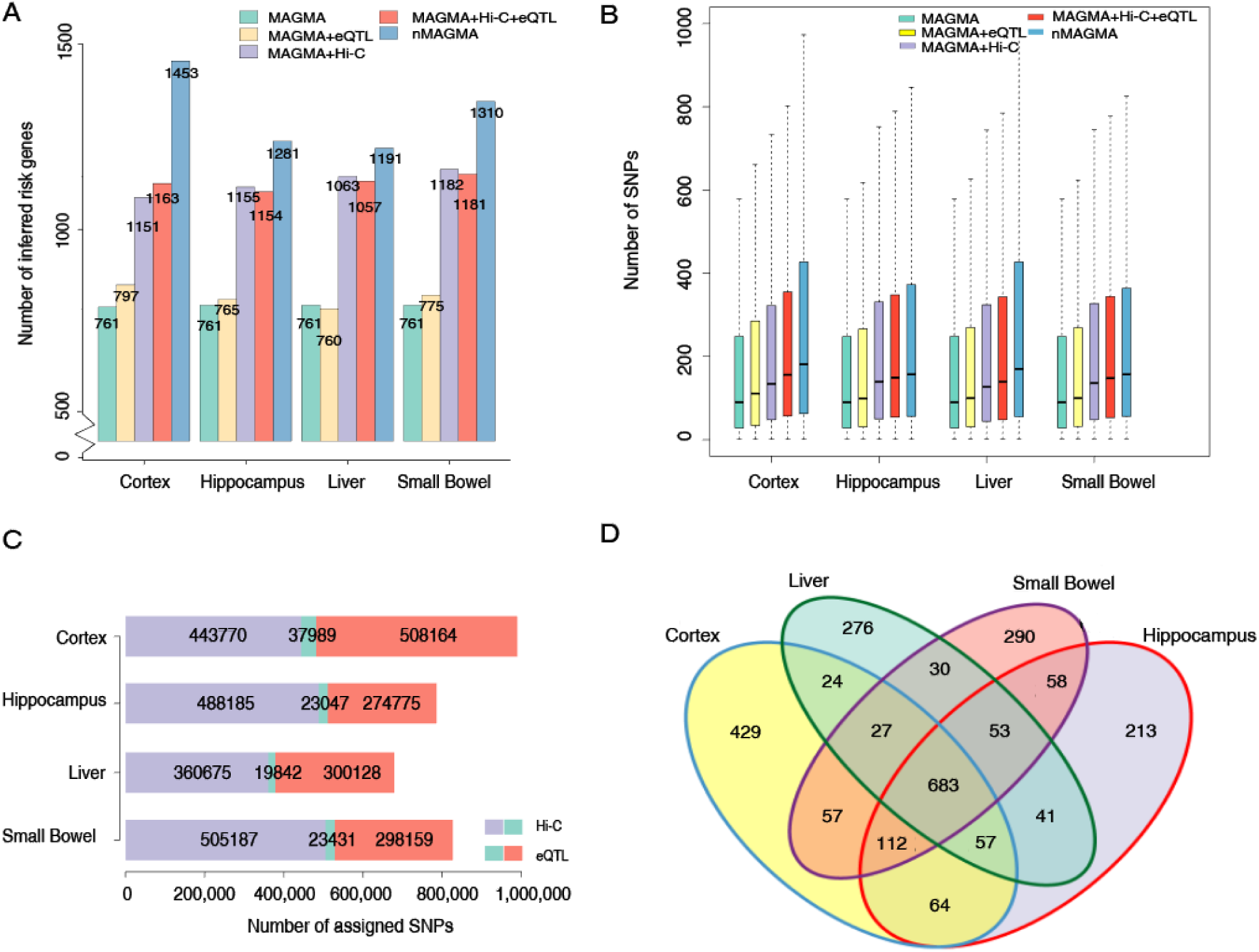
The gene-wise annotations of SNPs for SCZ by nMAGMA. **A**, The number of significant genes (p < 0.05, Bonferroni correction) detected by different strategies across four tissues. **B**, The median number of SNPs mapped to each gene by different strategies across four tissues. **C**, The number of SNPs that can be mapped to genes in four tissues through Hi-C and eQTL, respectively. **D**, The venn diagram of genes that can be detected by nMAGMA in four tissues.

### nMAGMA discovers more risk genes for SCZ

To evaluate the performance of nMAGMA, we compared it with MAGMA and H-MAGMA, a variant of MAGMA that also utilizes Hi-C information. Since the results by H-MAGMA are only available in Cortex, we compared nMAGMA against H-MAGMA in Cortex, while MAGMA has no tissue-specific results (same in all four tissues). Figure 3 shows the Venn diagrams of the results by the three approaches across four tissues. We can see that nMAGMA can detect more risk genes than both MAGMA and H-MAGMA, and has increased the identified risk genes by 57% compared to H-MAGMA and by 67% ∼ 100% compared to MAGMA (217% for a union of four tissues) (Figure 3A-D). nMAGMA has more consistent results with MAGMA, replicating about 89% ∼ 93% of the results by MAGMA (99% for union of four tissues results), while only 60% of MAGMA-detected genes can be replicated by H-MAGMA. 58% of the genes detected by H-MAGMA can be found by nMAGMA. From the results above, we can see that most of the results from MAGMA and H-MAGMA can be replicated by nMAGMA, which implies that the results from nMAGMA are confident and reliable.

**Figure 3:**
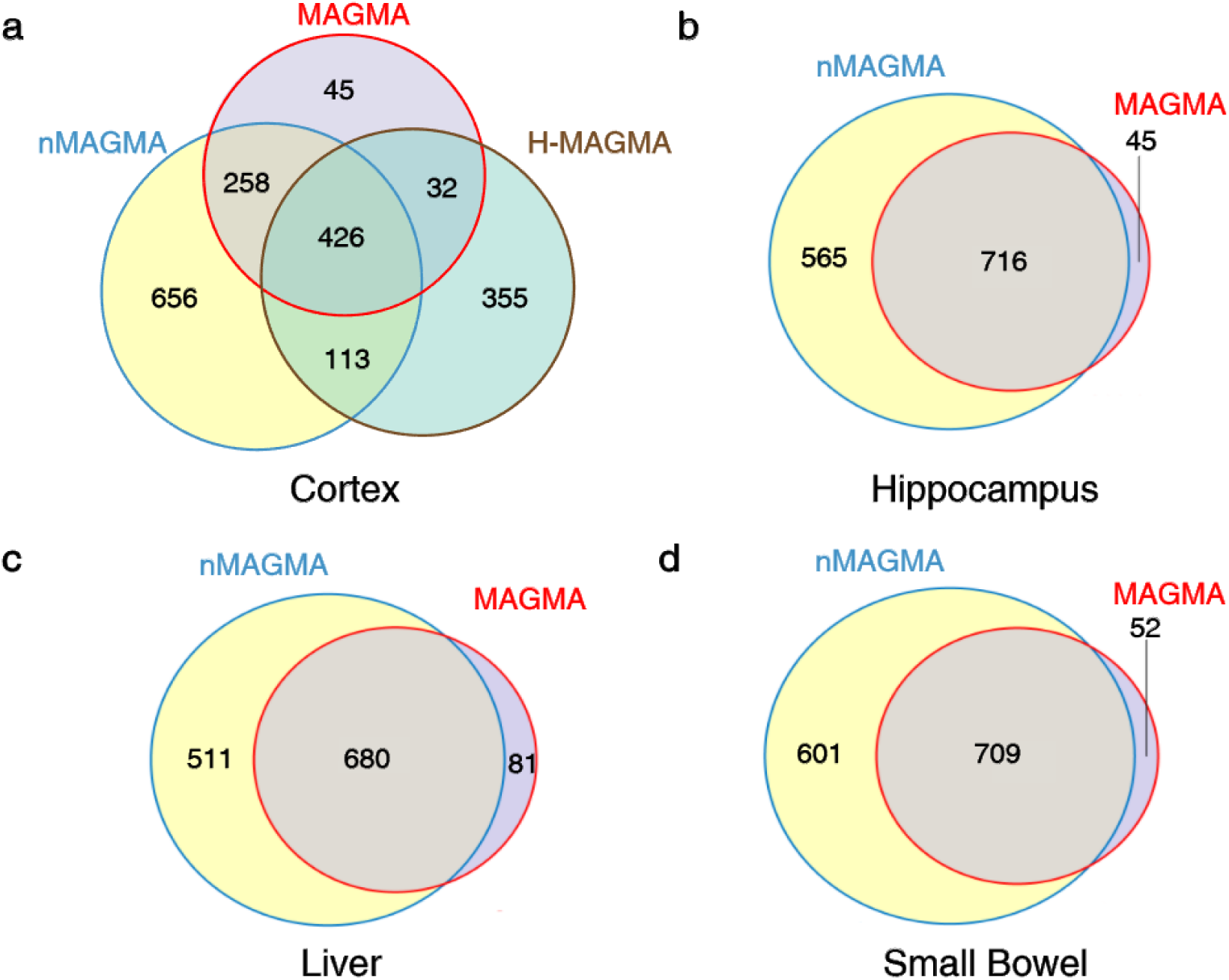
The venn diagram of risk genes inferred by nMAGMA, MAGMA and H-MAGMA.

To see whether nMAGMA can detect more confident risk genes, we validated the results by nMAGMA, H-MAGMA and MAGMA with known SCZ risk genes collected from public databases. For fair comparison, we compared the three approaches to see how much of their top predictions can be validated. Table 1 shows the results of the three tools with respect to their top 100, 500, 1500 and 2500 predictions. From the results, we can see that more of the results by nMAGMA can be validated with known SCZ risk genes in brain tissues, whereas the most confident risk genes can be detected based on their physical locations to the significant SNPs detected in GWAS, likely due to that many known SCZ genes are identified through previous GWAS studies. In addition, we validated our results with those by iRIGS, a useful approach to infer genes from risk loci of GWAS by integrating multi-omics data (e.g. Hi-C and de novo mutations) and gene networks derived from Gene Ontology [15]. A total of 116 risk loci were collected from the GWAS datasets we used (9 from [24] and 108 from [2] with one overlap), which were then used as input for iRIGS, resulting in 105 high-confidence risk genes identified by iRIGS. Among the 105 risk genes given by iRIGS, nMAGMA can successfully recover 49, 44, 41 and 45 genes in Cortex, Hippocampus, Liver, and Small Bowel, respectively. On the other hand, MAGMA can recover 41 iRIGS risk genes and H-MAGMA can also recover 41 iRIGS risk genes. The validation of the results by nMAGMA with known or other predicted risk genes imply that nMAGMA is reliable for gene-wise SNP annotations.

**Table 1:** Comparison of the results by MAGMA, H-MAGMA and nMAGMA, and the numbers of genes that can be validated with known SCZ risk genes.

| **Methods** | **Top 100** | **Top 500** | **Top 1500** | **Top 2500** | **iRIGS genes** |
| --- | --- | --- | --- | --- | --- |
| **MAGMA** | 43 | 144 | 303 | 425 | 47 |
| **H-MAGMA** | 35 | 128 | 290 | 425 | 47 |
| **nMAGMA** |  |  |  |  |  |
| Cortex | 37 | 153 | 319 | 455 | 49 |
| Hippocampus | 34 | 137 | 306 | 440 | 46 |
| Liver | 35 | 144 | 297 | 415 | 46 |
| Small Bowel | 34 | 145 | 310 | 433 | 48 |

By looking over the risk genes that can be detected by all three tools, we noticed that most of them have function known related to SCZ, e.g. the calcium channel and signaling genes (*CACNA1C, CACNB2*), the glutamatergic neurotransmission genes (*GRIN2A, GRM3*) and the neurogenesis genes (*SATB2, SOX2*). By examining the unique risk genes inferred by nMAGMA, we noticed that many of them have been reported related to SCZ. For instance, the gene *H2BC9* (Cortex: p = 5×10^−12^, Hippocampus: p = 7×10^−10^) was differentially expressed in lymphoblastoid cell lines of SCZ cases versus controls [25]. *H2BC9* is a member of the H2B histone family and is located in the extended major histocompatibility complex (xMHC) region, which is found to be the locus containing several most significant variants for SCZ in a GWAS study [26]. In addition to the brain tissues, Liver and Small Bowel have also been found involved in SCZ from our results. nMAGMA predicted *AKT1* (v-akt murine thymoma viral oncogene homolog 1) from Liver (p = 1.12×10^−6^) and *SYN2* (Synapsin II) from Small Bowel (p = 1.85×10^−11^) to be risk genes of SCZ. *AKT1* has been reported to be a SCZ risk gene related to neurocognition [27], and cognitive deficit is a typical symptom in SCZ [28]. The gene *SYN2* functions in synaptogenesis, and the dysfunction of synaptic transmission is known in the fundamental pathology of SCZ [29, 30]. The risk genes inferred above may confirm the involvement of non-brain tissues in SCZ [10, 21].

### nMAGMA identifies more meaningful gene-sets for SCZ

With the genes identified above by the three approaches, we next investigated the gene-sets enriched in those genes to see how the genes are involved in schizophrenia. The genes inferred by nMAGMA from four tissues were enriched in 32 (Cortex), 67 (Hippocampus), 32 (Liver) and 38 (Small Bowel) gene-sets, respectively. The genes inferred by MAGMA were enriched in 19 gene-sets while only 4 gene-sets were enriched with genes inferred by H-MAGMA. The gene-sets detected by MAGMA include 6 synaptic and neuronal gene-sets and 2 RNA-binding gene-sets, while the gene-sets detected by H-MAGMA have similar functions. It has been reported that the dysfunction of synaptic and neuronal genes is involved in SCZ and Bipolar disorders [31], and the copy number variation of RNA binding proteins like *RBFOX1* contributes to the pathology of SCZ and autism [32, 33].

We are more interested in the gene-sets detected by nMAGMA but not by MAGMA and H-MAGMA. Compared to the general synaptic functions enriched in the genes detected by MAGMA and H-MAGMA, the results by nMAGMA imply that both pre-synaptic and post-synaptic processes are involved in SCZ, which is consistent with previous findings that post-synaptic glutamatergic process and molecules regulating presynaptic transmitter release are both implicated in SCZ [34, 35]. Interestingly, nMAGMA identified many gene-sets related to CHD8 (chromodomain-helicase-DNA binding protein 8). CHD8 is a chromatin modifier which encodes the ATP-dependent chromatin-remodeling factors, and was found related to neurodevelopmental disorders like SCZ and autism by bothering gene expression and regulation in human brain [36-38]. It is noteworthy that the gene-sets associated with other non-psychiatric diseases (e.g. Breast cancer, Crohn’s disease and Liver cancer) were also found significant in nMAGMA, in accordance with previous findings that people with SCZ also suffer from these non-psychiatric disorders at the same time [39-42].

Table 2 shows the functional categories of the gene-sets detected by nMAGMA across four tissues. We noticed that the ATPase-related gene-sets were detected in Cortex but not in the other three tissues. ATPase like Na+/K+ ATPase-α1 is a potential modulator of glutamate ingestion in the brain, and has been reported to contribute to the pathophysiology of schizophrenia in Prefrontal Cortex [43]. Most of the gene-sets detected in both Cortex and Hippocampus have been found related to SCZ, such as those enriched in biological processes on synapse, oligodendrocyte and FMRP. Oligodendrocytes belongs to glial cells in the nervous system, and the dysfunction of oligodendrocytes has been frequently found associated with SCZ and bipolar disorders [44, 45]. The fragile-X mental retardation protein (FMRP) affects the development and maturation of synapses between neurons, and the loss of FMRP can inhibit neuronal translation and synaptic function, causing psychiatric symptoms including autistic and schizophrenic features [46, 47]. Among the two non-brain tissues, we noticed that the immune functions (e.g. T cell, B cell and Interferon) were extensively affected in Liver, which involves multiple cells like lymphocytes that are related to the immunoreaction processes [48]. This suggests that the dysfunction of the immune processes in Liver may be related to SCZ. What’s more, the metabolic processes (RNA, acid and amines metabolism) were found significant in Small Bowel, where the abnormal metabolic processes caused by unhealthy intestinal microbiota can influence brain chemistry and contributes to psychiatric disorders [49, 50]. The observation of SCZ-associated gene-sets in Liver and Small Bowel implies the association between non-brain tissues and psychiatric disorders.

**Table 2:** Biological processes underlying gene-sets detected by nMAGMA for schizophrenia across tissues.

| Function | Cortex | Hippocampus | Liver | Small Bowel |
| --- | --- | --- | --- | --- |
| Transcription & Regulation | ✓ | ✓ | ✓ | ✓ |
| ATPase | ✓ | - | - | - |
| Oligodendrocytes | ✓ | ✓ | - | - |
| Synaptic | ✓ | ✓ | - | ✓ |
| Neuronal related | ✓ | ✓ | ✓ | ✓ |
| Transmembrane movement of ions | ✓ | - | - | - |
| FMRP | ✓ | - | - | - |
| Cellular organization, cellular component, movement | ✓ | ✓ | ✓ | ✓ |
| RNA binding | - | ✓ | - | ✓ |
| Metabolic process | ✓ | ✓ |  | ✓ |
| Adipogenesis | - | - | - | ✓ |
| T cell | ✓ | ✓ | ✓ | - |
| B cell | - | - | ✓ | - |
| CALB1 | ✓ | ✓ | - | - |
| Protein targeting & transport | ✓ | ✓ | ✓ | ✓ |
| Interferon | ✓ | - | ✓ | - |
| Other diseases related | Alzheimers, Bipolar disorder, Crohn's disease, Bladder cancer | Alzheimers, Bipolar disorder, Myeloma, Thyroid cancer, Bladder cancer | Breast cancer, Liver cancer, Crohn's disease, Brain Glioma | Myeloma, Obesity |

### The network modules uncover biological processes implicated in SCZ

We further looked at the gene networks constructed by WGCNA, where two genes were linked if their TOM is no less than 0.15 (see Methods). We focused on the genes significant in nMAGMA. We are more interested in the module structures consist of these intensely connected genes, which were detected by Molecular Complex Detection (MCODE) [51] with default parameters, and only the modules with more than 10 genes were kept. At last, a total of 16 modules were detected across the four tissues as shown in Table 3. The functional terms enriched in the 16 modules were detected with the Database for Annotation, Visualization and Integrated Discovery (DAVID, version 6.8) [52], where only the most enriched term was shown for each module in Table 3 for clarity.

**Table 3:**
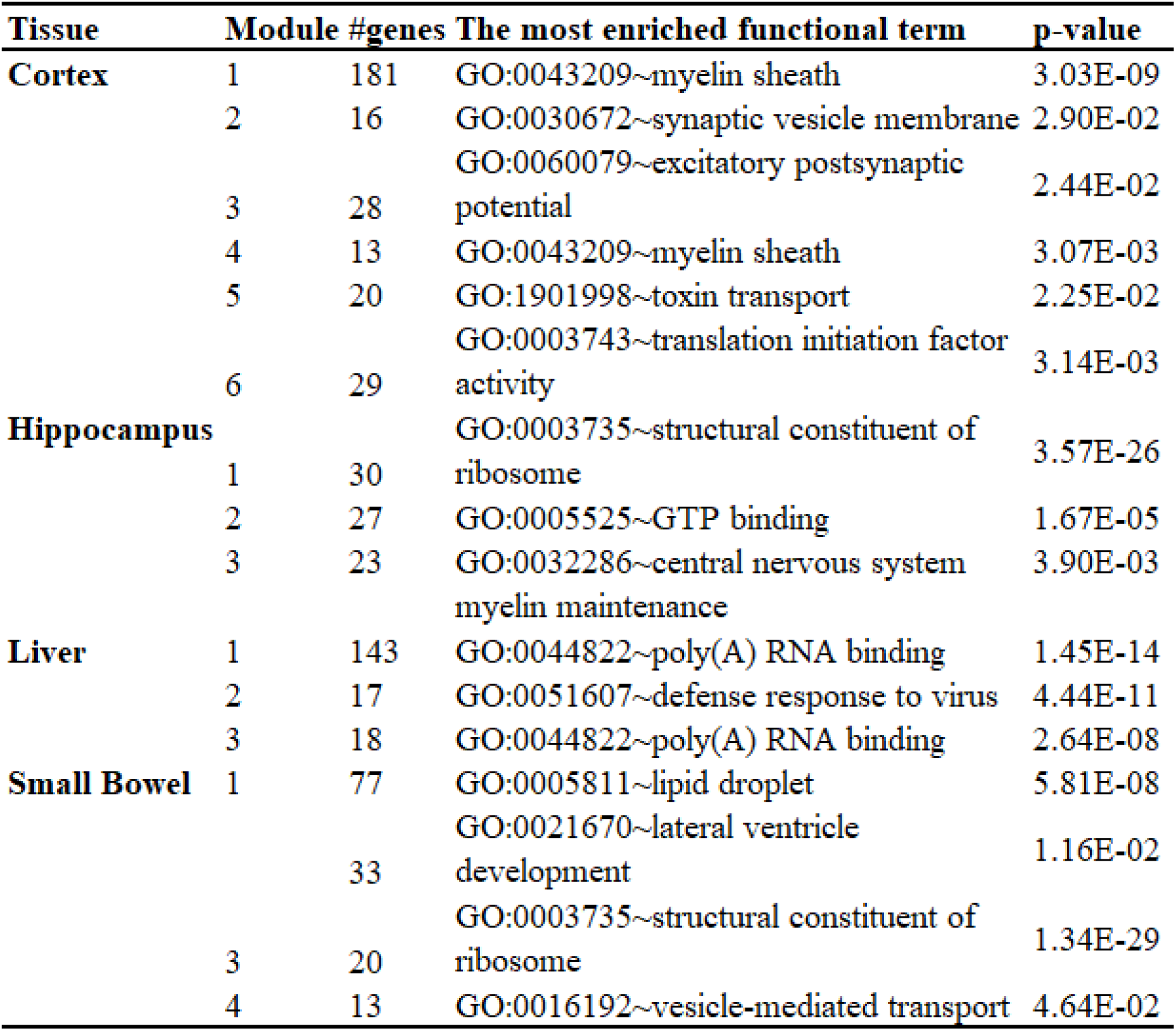
Details of the 16 modules detected by nMAGMA across four tissues, and their most enriched functional terms.

Figure 4 shows the 6 modules detected in Cortex, which was visualized with Cytoscape (Version 3.7.2) [53]. Module 1 contains 181 genes, and was enriched with SCZ risk genes (including 49 risk genes). Module 1 was enriched in myelin sheath (p=3.03×10^−9^), which is a membrane surrounding the axons of neurons and is the outgrowth of glial cells. Abnormality of myelin has been reported to be causally involved in SCZ in the Prefrontal Cortex [54], where several myelin-related genes were also found reduced expression in SCZ patients compared to controls [55]. What’s more, the gene *RTN4* in Module 1 which encodes myelin-related proteins, was detected differential expression in the Cortex of SCZ by inhibiting neurodevelopment in the previous research [56]. The myelin-related function was also enriched in module 4 of Cortex and module 3 of Hippocampus.

**Figure 4:**
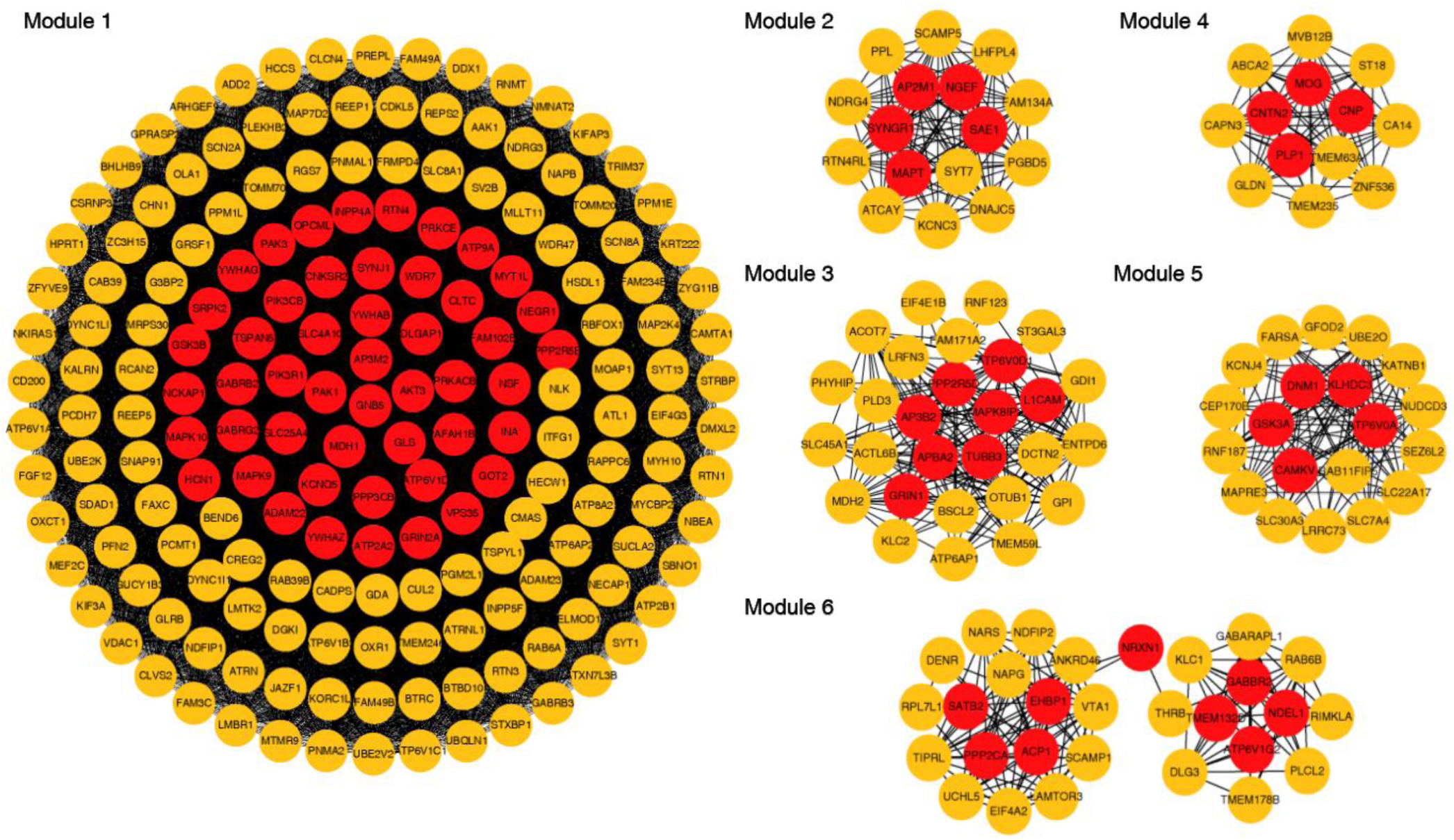
Visualization of modules in Cortex. The red node represents a known SCZ risk gene.

The functional term of structural constituent of ribosome was found most enriched in Hippocampus (module 1) and Small Bowel (module 3), as well as an immune-related term of defense response to virus in Liver. These agree with former studies on functional enrichment of differentially expressed genes or long noncoding RNA in SCZ, where pathways associated with ribosome and immune response were also enriched [57, 58]. Another, the functional component of lipid droplet in Small Bowel (p=5.81×10^−8^, module 1) is a main organelle that has important roles in regulating lipid metabolism [59]. Slight hurt of lipid metabolism in SCZ cases under risperidone monotherapy (a first-choice for SCZ treatment) was found before [60]. Altogether, our analysis validates that network metrics can help to identify new groups of interconnected genes with known roles in SCZ and reveal hidden enrichments in both brain and non-brain tissues.

## Discussion

In this paper, we provide a method leveraging multidimensional resources to extract comprehensive biological information from GWAS. The challenge is to incorporate tissue-specific gene networks into annotation, which have been proven helpful for prioritizing candidate disease genes that may otherwise be ignored by conventional approaches. Instead of roughly computing correlations among genes, nMAGMA specifies the gene regulatory relationships from TOM matrix which is more explicit and biologically meaningful to promote inference accuracy. To cover more genomic signals with high confidence, Hi-C and eQTL identified SNP-gene relationships were also taken into account in nMAGMA. Unlike many other studies that explore the expression profiles of selected significant genes, nMAGMA directly incorporates the expression data into the upstream analysis, making the downstream analysis more reliable.

Empowered by the integration of gene networks and multi-omics information, nMAGMA shows greater ability in detecting risk signatures for SCZ compared with MAGMA and H-MAGMA. Featuring tissue-specific results, nMAGMA also contributes to elucidating the underpinnings of the tissue-specific origins of diseases. Strong evidence of the latent role of non-brain tissues also presents in SCZ in our results, while non-brain tissues are less discussed in previous studies on psychiatric diseases. After characterizing the topology and sub-network structures of nMAGMA-detected genes, we show that network information can be used to uncover hidden disease-related enrichments.

Though results of nMAGMA are excellent in many aspects, there are still some potential directions for further expansion of our work. First, we only focused on four relevant tissues in this work. nMAGMA can be extended to other tissues by using other functional genomic resources such as BrainVar [79], for more precise tissue matching between multidimensional data. Second, more gene interaction resources can be utilized for the gene-wise annotation. Other types of networks, for example, the Protein-Protein Interaction networks [61] and the Disease Gene Networks [62], may be further incorporated. Ready-made genes interaction relations can also be obtained from international and multidisciplinary databases like BIND (http://www.bind.ca). Third, as the impact of each SNP on its target gene may not be the same, additional information can be added to give different weights to different SNP-gene relationships. For example, predicted scores to prioritize “pathogenic” SNPs, which combine functional information (e.g. variant impact on transcripts), positional information (e.g. intronic), sequence conservation information, etc., can be used as weights in gene analysis. nMAGMA is designed and implemented in a way easy to expand, and very flexible for incorporating new information.

Altogether, nMAGMA provides a strategy for using network information to select potentially risk genes and identify relevant biological function. Example of results show that nMAGMA may help improve our understanding of the complex genetic background governing the susceptibility to complex diseases such as SCZ. We hope this method can be of value for deciphering disease mechanisms and selecting therapeutic drug candidates, and that it can provide a basis for more innovative methods in this area.

## Materials and Methods

### GWAS data

The two SCZ GWAS summary statistics datasets were downloaded from Psychiatric Genomics Consortium (https://www.med.unc.edu/pgc/download-results/scz/). These two datasets only contain samples of European ancestry, where dataset1 contains 13,833 SCZ cases and 18,310 controls across 9,898,079 SNPs (from PGC1 + Sweden [24]), and dataset2 contains 35,476 SCZ cases and 46,839 controls across 15,358,497 SNPs (from PGC2 [2]). For more details about the datasets please refer to the original publications [2, 24].

### eQTL and gene expression data

The significant SNP-gene pairs (FDR<0.05) were retrieved from Genotype-Tissue Expression (GTEx) (version 8) [63, 64], where the SNP-gene associations were obtained from cis-eQTL analysis with linear regression of gene expression against SNP genotypes. Specifically, the eQTLs and gene expression (TPM) data from the four tissues related to SCZ were considered here, including Cortex, Hippocampus, Liver and Small Bowel (Small Intestine Terminal Ileum). All the gene IDs were transformed to gene names in Ensemble (GRCH37 v87) [65], and all variant IDs were transformed to RS IDs in dbSNP (version 151) with a table provided by GTEx.

### Hi-C data

The chromatin interaction profiles based on Hi-C data were collected from 3D-genome Interaction Viewer and database [66] for four tissues, i.e. adult Dorsolateral Prefrontal Cortex (DLPFC, fuzzily matched with Cortex), Hippocampus, Liver, and Small Bowel. The chromatin interaction regions overlapped with the regulatory elements (REs) defined in ENCODE [67] and Ensembl and with protein-coding genes from Ensemble (GRCH37 v87) were kept for further analysis. An interaction of an RE and a gene, based on the chromatin interaction profiles, indicates that the RE has the potential to affect the expression level of the gene.

### SCZ risk genes

To verify the risk genes inferred for SCZ, a set of genes associated with SCZ were collected from public resources. The gene list contains about 2400 candidate SCZ risk genes collected from public databases: SZGR2 [68], GAD [69], OMIM [70] and SzGene [71]. SZGR2 genes contain those collected from genetics studies (such as genes harboring GWAS loci), transcriptome studies (such as differentially expressed genes) and epigenetics studies (such as differentially methylated genes), while GAD, OMIM and SzGene genes are collected from thousands of genetic association studies.

### Gene sets for functional analysis

A total of 17224 gene-sets were used for gene-set analysis, including those used previously [72] and agnostic gene-sets downloaded from the Molecular Signatures Database (MSigDB, version 7.1) [73, 74], where only the MSigDB gene-sets with more than ten and less than 1500 genes were considered. The details of the gene-sets can be found in Supplementary Table S1.

### Network-enhanced MAGMA (nMAGMA)

We only took protein-coding genes for further analysis, where the protein-coding genes and the coordinates of transcription regions were extracted from Ensemble (GRCH37 v87). In nMAGMA, we inferred risk genes from SNPs with the following steps: Firstly, the SNPs were mapped to genes and REs using MAGMA based on their physical locations, where a 0kb-window was adopted for the European population; Secondly, the SNPs were assigned to genes via SNP-to-RE annotations and RE-gene regulatory pairs derived from Hi-C-based chromatin interaction profiles; Thirdly, the SNPs were assigned to genes based on significant eQTL results from GTEx; Finally, the topological overlap matrix (TOM) was calculated with WGCNA for genes based on their expression profiles from GTEx, where a pair of genes with a TOM ≥ 0.15 were regarded as strongly interconnected as described previously [75]. In this way, the lists of genes inferred from the first three steps can be further extended, where a gene that has a TOM ≥ 0.15 with any gene inferred from the first three steps will be included in analysis. Finally, we got a final SNP-gene mapping annotation file, with which we can perform gene analysis to determine the association of a certain gene with SCZ, where the linkage disequilibrium (LD) information was taken from the 1000 Genomes Project EUR panels (Phase 3) for estimating the LD between SNPs in a gene. Then the meta-analysis of the two GWAS summary datasets was performed, followed the gene-set analysis.

### Comparison with MAGMA and H-MAGMA

To show the performance of nMAGMA, we compared it with MAGMA and H-MAGMA to see how much of their detected significant genes (p-value < 0.05 after Bonferroni correction) and gene-sets (p-value < 5×10^−6^) can be validated with known SCZ risk genes. The annotation file of H-MAGMA (Adult_brain.genes.annot) was downloaded from https://github.com/thewonlab/H-MAGMA, where only protein-coding genes were kept for a fair comparison. The potential SCZ risk genes and gene-sets were subsequently inferred with the EUR samples we used for nMAGMA. Since no other tissues except for DLPFC are considered for H-MAGMA, we only compared nMAGMA with H-MAGMA for Cortex. We tested the overlap between nominally significant genes detected by the three methods against the known SCZ risk genes. Significant GO gene-sets with low levels (≤ 5) or having a ‘is_a’ or ‘part_of’ relationship with another significant GO term were removed.

### Availability

Codes and data used in this work are freely available at https://github.com/sldrcyang/nMAGMA.

## Acknowledgments

We deeply appreciate the researchers comprise the Psychiatric Genomics Consortium (PGC), and the hundreds of thousands of contributors who have provided the data.

## References

1. Pardinas AF, Holmans P, Pocklington AJ et al. Common schizophrenia alleles are enriched in mutation-intolerant genes and in regions under strong background selection, Nat Genet 2018;50:381–389.

2. Schizophrenia Working Group of the Psychiatric Genomics C. Biological insights from 108 schizophrenia-associated genetic loci, Nature 2014;511:421–427.

3. Cooper GM, Goode DL, Ng SB et al. Single-nucleotide evolutionary constraint scores highlight disease-causing mutations 2010;7:250–251.

4. Cooper GM, Shendure J. Needles in stacks of needles: finding disease-causal variants in a wealth of genomic data, Nature Reviews Genetics 2011;12:628–640.

5. Ward LD, Kellis M. Interpreting noncoding genetic variation in complex traits and human disease, Nature Biotechnology 2012;30:1095–1106.

6. de Leeuw CA, Mooij JM, Heskes T et al. MAGMA: generalized gene-set analysis of GWAS data, PLoS Comput Biol 2015;11:e1004219.

7. de Leeuw CA, Neale BM, Heskes T et al. The statistical properties of gene-set analysis, Nat Rev Genet 2016;17:353–364.

8. Holmans P. Statistical methods for pathway analysis of genome-wide data for association with complex genetic traits, Adv Genet 2010;72:141–179.

9. Chen JQ, Tian WD. Explaining the disease phenotype of intergenic SNP through predicted long range regulation, Nucleic Acids Research 2016;44:8641–8654.

10. Gamazon ER, Zwinderman AH, Cox NJ et al. Multi-tissue transcriptome analyses identify genetic mechanisms underlying neuropsychiatric traits, Nat Genet 2019;51:933–940.

11. Yin Z, Guo B, Mi Z et al. Gene Saturation: An Approach to Assess Exploration Stage of Gene Interaction Networks, Sci Rep 2019;9:5017.

12. Sey NYA, Hu B, Mah W et al. A computational tool (H-MAGMA) for improved prediction of brain-disorder risk genes by incorporating brain chromatin interaction profiles, Nat Neurosci 2020;23:583–593.

13. Gerring ZF, Gamazon ER, Derks EM et al. A gene co-expression network-based analysis of multiple brain tissues reveals novel genes and molecular pathways underlying major depression, PLoS Genet 2019;15:e1008245.

14. Radulescu E, Jaffe AE, Straub RE et al. Identification and prioritization of gene sets associated with schizophrenia risk by co-expression network analysis in human brain, Mol Psychiatry 2020;25:791–804.

15. Wang Q, Chen R, Cheng F et al. A Bayesian framework that integrates multiomics data and gene networks predicts risk genes from schizophrenia GWAS data, Nature Neuroscience 2019;22:691–699.

16. Ravasz E, Somera AL, Mongru DA et al. Hierarchical organization of modularity in metabolic networks, Science 2002;297:1551–1555.

17. Zhang B, Horvath S. A general framework for weighted gene co-expression network analysis, Stat Appl Genet Mol Biol 2005;4:Article17.

18. Langfelder P, Horvath S. WGCNA: an R package for weighted correlation network analysis, BMC Bioinformatics 2008;9:559.

19. Jaffe AE, Straub RE, Shin JH et al. Developmental and genetic regulation of the human cortex transcriptome illuminate schizophrenia pathogenesis, Nat Neurosci 2018;21:1117–1125.

20. Gallinat J, McMahon K, Kuhn S et al. Cross-sectional Study of Glutamate in the Anterior Cingulate and Hippocampus in Schizophrenia, Schizophr Bull 2016;42:425–433.

21. Kroll J. Schizophrenia and liver dysfunction, Med Hypotheses 2001;56:634–636.

22. Dong J, Horvath S. Understanding network concepts in modules, BMC Syst Biol 2007;1:24.

23. Rao SSP, Huntley MH, Durand NC et al. A 3D map of the human genome at kilobase resolution reveals principles of chromatin looping, Cell 2014;159:1665–1680.

24. Ripke S, O’Dushlaine C, Chambert K et al. Genome-wide association analysis identifies 13 new risk loci for schizophrenia, Nat Genet 2013;45:1150–1159.

25. Sanders AR, Goring HH, Duan J et al. Transcriptome study of differential expression in schizophrenia, Hum Mol Genet 2013;22:5001–5014.

26. Shi J, Levinson DF, Duan J et al. Common variants on chromosome 6p22.1 are associated with schizophrenia, Nature 2009;460:753–757.

27. Chow TJ, Tee SF, Yong H et al. Genetic Association of TCF4 and AKT1 Gene Variants with the Age at Onset of Schizophrenia, Neuropsychobiology 2016;73:233–240.

28. Gonzalez-Burgos G, Fish KN, Lewis DA. GABA neuron alterations, cortical circuit dysfunction and cognitive deficits in schizophrenia, Neural Plast 2011;2011:723184.

29. Hilfiker S, Pieribone VA, Czernik AJ et al. Synapsins as regulators of neurotransmitter release, Philosophical Transactions of the Royal Society of London Series B-Biological Sciences 1999;354:269–279.

30. Frankle WG, Lerma J, Laruelle M. The synaptic hypothesis of schizophrenia, Neuron 2003;39:205–216.

31. Bipolar D, Schizophrenia Working Group of the Psychiatric Genomics Consortium. Electronic address drve, Bipolar D et al. Genomic Dissection of Bipolar Disorder and Schizophrenia, Including 28 Subphenotypes, Cell 2018;173:1705–1715 e1716.

32. Weyn-Vanhentenryck SM, Mele A, Yan Q et al. HITS-CLIP and integrative modeling define the Rbfox splicing-regulatory network linked to brain development and autism, Cell Rep 2014;6:1139–1152.

33. Xu B, Roos JL, Levy S et al. Strong association of de novo copy number mutations with sporadic schizophrenia 2008;40:880–885.

34. Zhao Z, Xu J, Chen J et al. Transcriptome sequencing and genome-wide association analyses reveal lysosomal function and actin cytoskeleton remodeling in schizophrenia and bipolar disorder, Molecular Psychiatry 2015;20:563–572.

35. Hall LS, Medway CW, Pain O et al. A transcriptome-wide association study implicates specific pre- and post-synaptic abnormalities in schizophrenia, Hum Mol Genet 2020;29:159–167.

36. Cotney J, Muhle RA, Sanders SJ et al. The autism-associated chromatin modifier CHD8 regulates other autism risk genes during human neurodevelopment, Nat Commun 2015;6:6404.

37. Thompson BA, Tremblay V, Lin G et al. CHD8 is an ATP-dependent chromatin remodeling factor that regulates beta-catenin target genes, Molecular and Cellular Biology 2008;28:3894–3904.

38. Gonzalez-Mantilla AJ, Moreno-De-Luca A, Ledbetter DH et al. A Cross-Disorder Method to Identify Novel Candidate Genes for Developmental Brain Disorders, JAMA Psychiatry 2016;73:275–283.

39. Izci F, Ilgun AS, Findikli E et al. Psychiatric Symptoms and Psychosocial Problems in Patients with Breast Cancer, Journal of Breast Health 2016;12:94–101.

40. Fuller BE, Rodriguez VL, Linke A et al. Prevalence of liver disease in veterans with bipolar disorder or schizophrenia, General Hospital Psychiatry 2011;33:232–237.

41. West J, Logan RF, Hubbard RB et al. Risk of schizophrenia in people with coeliac disease, ulcerative colitis and Crohn’s disease: a general population-based study, Aliment Pharmacol Ther 2006;23:71–74.

42. Barnett AH, Mackin P, Chaudhry I et al. Minimising metabolic and cardiovascular risk in schizophrenia: diabetes, obesity and dyslipidaemia 2007;21:357–373.

43. Corti C, Xuereb JH, Crepaldi L et al. Altered levels of glutamatergic receptors and Na+/K+ ATPase-alpha1 in the prefrontal cortex of subjects with schizophrenia, Schizophr Res 2011;128:7–14.

44. Tkachev D, Mimmack ML, Ryan MM et al. Oligodendrocyte dysfunction in schizophrenia and bipolar disorder, Lancet 2003;362:798–805.

45. Goudriaan A, de Leeuw C, Ripke S et al. Specific glial functions contribute to schizophrenia susceptibility, Schizophr Bull 2014;40:925–935.

46. Darnell JC, Van Driesche SJ, Zhang C et al. FMRP stalls ribosomal translocation on mRNAs linked to synaptic function and autism, Cell 2011;146:247–261.

47. Fatemi SH, Folsom TD. GABA receptor subunit distribution and FMRP–mGluR5 signaling abnormalities in the cerebellum of subjects with schizophrenia, mood disorders, and autism, Schizophrenia Research 2015;167:42–56.

48. Nemeth E, Baird AW, O’Farrelly C. Microanatomy of the liver immune system, Semin Immunopathol 2009;31:333–343.

49. Bercik P, Denou E, Collins J et al. The intestinal microbiota affect central levels of brain-derived neurotropic factor and behavior in mice, Gastroenterology 2011;141:599–609, 609 e591-593.

50. Ojeda P, Bobe A, Dolan K et al. Nutritional modulation of gut microbiota - the impact on metabolic disease pathophysiology, J Nutr Biochem 2016;28:191–200.

51. Bader GD, Hogue CW. An automated method for finding molecular complexes in large protein interaction networks, BMC Bioinformatics 2003;4:2.

52. Huang DW, Sherman BT, Lempicki RA. Systematic and integrative analysis of large gene lists using DAVID bioinformatics resources, Nature Protocols 2009;4:44–57.

53. Shannon P, Markiel A, Ozier O et al. Cytoscape: a software environment for integrated models of biomolecular interaction networks, Genome Res 2003;13:2498–2504.

54. Regenold WT, Phatak P, Marano CM et al. Myelin staining of deep white matter in the dorsolateral prefrontal cortex in schizophrenia, bipolar disorder, and unipolar major depression, Psychiatry Res 2007;151:179–188.

55. Hof PR, Haroutunian V, Friedrich VL, Jr. et al. Loss and altered spatial distribution of oligodendrocytes in the superior frontal gyrus in schizophrenia, Biol Psychiatry 2003;53:1075–1085.

56. Novak G, Kim D, Seeman P et al. Schizophrenia and Nogo: elevated mRNA in cortex, and high prevalence of a homozygous CAA insert, Brain Res Mol Brain Res 2002;107:183–189.

57. Tian T, Wei Z, Chang X et al. The Long Noncoding RNA Landscape in Amygdala Tissues from Schizophrenia Patients, EBioMedicine 2018;34:171–181.

58. Chang X, Liu Y, Hahn CG et al. RNA-seq analysis of amygdala tissue reveals characteristic expression profiles in schizophrenia, Transl Psychiatry 2017;7:e1203.

59. Vallochi AL, Teixeira L, Oliveira KDS et al. Lipid Droplet, a Key Player in Host-Parasite Interactions, Front Immunol 2018;9:1022.

60. Murashita, MariInoue, Kusumi T et al. Glucose and lipid metabolism of long-term risperidone monotherapy in patients with schizophrenia, Psychiatry Clinical Neuroences 2010;61:54–58.

61. Rual JF, Venkatesan K, Hao T et al. Towards a proteome-scale map of the human protein-protein interaction network, Nature 2005;437:1173–1178.

62. Goh KI, Cusick ME, Valle D et al. The human disease network, Proc Natl Acad Sci U S A 2007;104:8685–8690.

63. Consortium GT, Laboratory DA, Coordinating Center-Analysis Working G et al. Genetic effects on gene expression across human tissues, Nature 2017;550:204–213.

64. Consortium GT. Human genomics. The Genotype-Tissue Expression (GTEx) pilot analysis: multitissue gene regulation in humans, Science 2015;348:648–660.

65. Hunt SE, McLaren W, Gil L et al. Ensembl variation resources, Database (Oxford) 2018;2018.

66. Yang D, Jang I, Choi J et al. 3DIV: A 3D-genome Interaction Viewer and database, Nucleic Acids Res 2018;46:D52–D57.

67. Consortium EP. An integrated encyclopedia of DNA elements in the human genome, Nature 2012;489:57–74.

68. Jia P, Sun J, Guo AY et al. SZGR: a comprehensive schizophrenia gene resource, Mol Psychiatry 2010;15:453–462.

69. Becker KG, Barnes KC, Bright TJ et al. The genetic association database, Nat Genet 2004;36:431–432.

70. Amberger JS, Bocchini CA, Schiettecatte F et al. OMIM.org: Online Mendelian Inheritance in Man (OMIM(R)), an online catalog of human genes and genetic disorders, Nucleic Acids Res 2015;43:D789–798.

71. Allen NC, Bagade S, McQueen MB et al. Systematic meta-analyses and field synopsis of genetic association studies in schizophrenia: the SzGene database, Nat Genet 2008;40:827–834.

72. Lam M, Chen CY, Li Z et al. Comparative genetic architectures of schizophrenia in East Asian and European populations, Nat Genet 2019;51:1670–1678.

73. Subramanian A, Tamayo P, Mootha VK et al. Gene set enrichment analysis: a knowledge-based approach for interpreting genome-wide expression profiles, Proc Natl Acad Sci U S A 2005;102:15545–15550.

74. Liberzon A, Subramanian A, Pinchback R et al. Molecular signatures database (MSigDB) 3.0, Bioinformatics 2011;27:1739–1740.

75. Farber CR. Systems-level analysis of genome-wide association data, G3 (Bethesda) 2013;3:119–129.

